# Developmental transitions coordinate assembly of the *Coxiella burnetii* Dot/Icm type IV secretion system

**DOI:** 10.1101/2022.05.29.493899

**Authors:** Donghyun Park, Samuel Steiner, Meng Shao, Craig R. Roy, Jun Liu

## Abstract

*Coxiella burnetii* is an obligate intracellular bacterial pathogen that has evolved a unique bi-phasic developmental cycle. The infectious form of *C. burnetii* is the dormant small cell variant (SCV) that transitions to a metabolically active large cell variant (LCV) that replicates inside the lysosome-derived host vacuole. A Dot/Icm type IV secretion system (T4SS), which can deliver over 100 effector proteins to host cells, is essential for the biogenesis of the vacuole and for intracellular replication. How the distinct *C. burnetii* life cycle impacts the assembly and function of the Dot/Icm T4SS has remained unknown. Here, we combine advanced cryo-focused ion beam (cryo-FIB) milling and cryo-electron tomography (cryo-ET) imaging to visualize all developmental transitions and assembly of the Dot/Icm T4SS. Importantly, assembled Dot/Icm T4SSs were not present in the infectious SCV. The appearance of an assembled Dot/Icm machine correlated with the transition of the SCV to the LCV intracellularly. Furthermore, temporal characterization of *C. burnetii* morphological changes revealed regions of the inner membrane that invaginated to form tightly packed stacks during the LCV to SCV transition at late stages of infection, which could enable the SCV to LCV transition that occurs upon infection of a new host cell. Overall, these data establish how *C. burnetii* developmental transitions control critical bacterial processes to promote intracellular replication and transmission.

## INTRODUCTION

*Coxiella burnetii* (*C. burnetii*) is an obligate intracellular pathogen and the causative agent of the human disease Q fever^1,2^. Through the dominant mode of transmission, bacteria are shed by *C. burnetii*-infected animals as environmentally resistant spore-like particles that can be aerosolized and inhaled by a new host. With an infectious dose of fewer than 10 bacteria, *C. burnetii* constitutes one of the most infectious and transmissive bacterial pathogens^3^. Upon infection of a new host cell, the infectious and dormant spore-like bacterium called the small cell variant (SCV) differentiates into the metabolically active large cell variant (LCV), which is the replicative form of *C. burnetii*. Acidification of *Coxiella*-containing vacuoles (CCVs) by fusion with lysosomes triggers the transition of the SCV to the LCV^4-7^, and also serves as an environmental signal that correlates with the activation of the *C. burnetii* Dot/Icm type IV secretion system (T4SS) and the translocation of an arsenal of over 100 different effector proteins by this secretion system^8^. A functional Dot/Icm machine and several translocated effector proteins are thus essential for *C. burnetii* intracellular replication^9-12^.

T4SSs are multiprotein nanomachines that span the cell membranes of Gram-negative and Gram-positive bacteria^13-16^ and are major virulence determinants involved in pathogenesis and antibiotic resistance^17^. Phylogenetically, T4SSs can be divided into two distinct subtypes: type IVA and type IVB^18,19^. The assembled IVA machines typically have 8-12 different protein subunits, whereas, IVB systems show greater complexity and are composed of more than 20 different protein subunits. The Dot/Icm system represents a prototypical IVB machine and is encoded by the intracellular pathogens *Legionella pneumophila, Rickettsiella grylli*, and *C. burnetii*^20^. The *L. pneumophila* Dot/Icm system has been extensively studied and serves as a model system for understanding the architecture of IVB machines. Recent single-particle cryo-electron microscopy (cryo-EM) analysis of purified Dot/Icm machines revealed near-atomic resolution structures of the outer membrane complex and periplasmic ring and complex symmetry mismatches^21,22^. Cryo-electron tomography (cryo-ET) was deployed not only to characterize the intact Dot/Icm machine in *L. pneumophila*^23-26^ but also reveal unique subcomplexes likely involved in the assembly of the Dot/Icm machine^26^. Cryo-ET studies also identified an inner membrane channel that undergoes a conformational switch upon association with the cytoplasmic ATPases DotO and DotB beneath the inner membrane complex. This channel may mediate the passage of T4SS substrates across the inner membrane^24,26^.

Here, we combined multiple cryo-ET techniques to visualize *C. burnetii* variants at high resolution to determine how the SCV-to-LCV transition impacts bacterial cell biology and whether assembly of the Dot/Icm machine is subject to developmental regulation. Overall, these data establish a link between the SCV-to-LCV transition and assembly of the Dot/Icm system.

## RESULTS

### Visualization of intracellular *C. burnetii* with a cellular cryo-ET pipeline

*C. burnetii* has evolved unique strategies to evade host immunity and replicate inside host cells. *Coxiella*-containing vacuoles (CCVs) mature along the endocytic pathway of the host cell. The gradual acidification of the vacuole promotes the conversion of the infectious small cell variant (SCV) into the metabolically active large cell variant (LCV)^4,6,7^. This transition initiates *C. burnetii* replication inside the CCV. Upon completion of the intracellular life cycle, the LCV must transition back to the infectious SCV before bacteria are released from the infected cell (**Fig. 1a**). Although SCVs and LCVs have been visualized using transmission electron microscopy (TEM) of chemically fixed and resin-embedded specimens^27-29^, their structural details have been limited by this technique and sample preparation. To obtain a better understanding of the native spatial and temporal dynamics of the CCV and host environment, we visualized COS-7 cells infected with a *C. burnetii* strain constitutively producing the mCherry protein by combining advanced imaging techniques, including cryo-fluorescence microscopy, cryo-focused ion beam (cryo-FIB) milling, and cryo-ET imaging (**Supplementary Video 1**).

**Figure 1.**
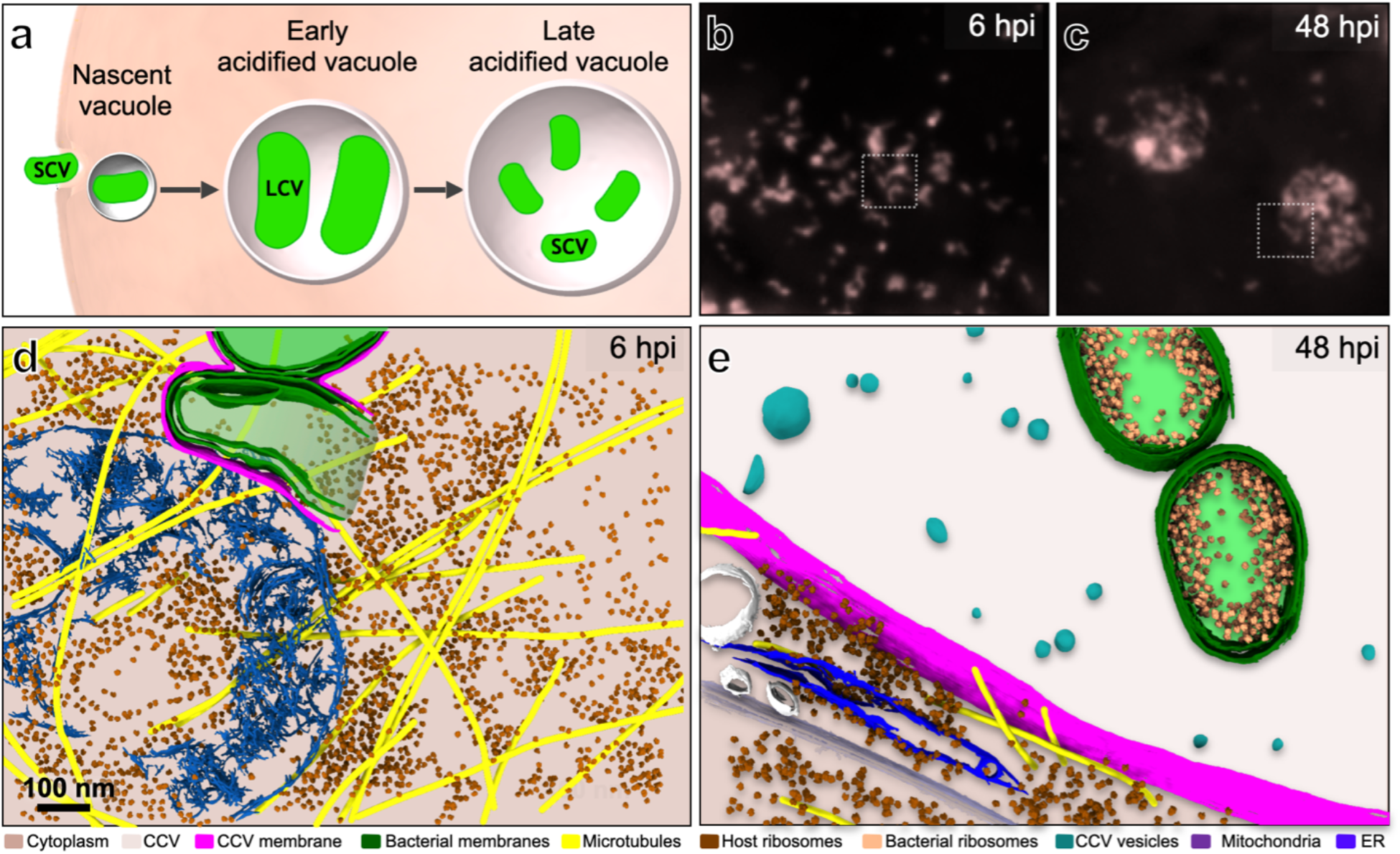
Visualization of the life cycle of the intracellular pathogen *C. burnetii* by combining cryo-fluorescence microscopy, FIB-milling, and cryo-ET. **(a)** A simple model of the life cycle of *C. burnetii* inside a host cell. **(b-c)** Cryo-fluorescence images of an mCherry-expressing *C. burnetii* strain show *Coxiella*-containing vacuoles (CCVs) at 6 hpi and 48 hpi, respectively. Areas in white squares were selected for cryo-FIB milling. **(d-e)** 3D renderings of the tomograms collected from thin lamellae generated by FIB-milling in the boxed areas in panels **b** and **c**, respectively.

Cryo-EM grids of infected COS-7 cells were plunge frozen at 6 and 48 hours post-infection (hpi), respectively (**Extended Data Fig. 1**). To identify host cells containing replicating bacteria inside a CCV, the vitrified samples were imaged on a cryo-fluorescence microscope to detect mCherry fluorescence. Consistent with previous studies, fluorescent objects, each representing an individual CCV, were observed at 6 hpi (**Fig. 1b**). At 48 hpi, fewer but larger CCVs occupied each cell, a change caused by homotypic fusion of independently derived CCVs contained in the same host cell^30^ (**Fig. 1c)**.

Location information of individual CCVs obtained by cryo-fluorescence microscopy was then used to prepare thin cryo-lamellae containing CCVs using the cryo-FIB method (**Extended Data Fig. 1**). At 6 hpi, cryo-ET imaging of the cryo-lamellae showed that CCVs with one or a few bacterial cells were tightly surrounded by the vacuolar membrane, with an average membrane spacing of ∼13.7 nm (**Fig. 1d, Extended Data Fig. 2a, b, e**). At 48 hpi, multiple bacteria appeared as dispersed objects within a spacious CCV (**Fig. 1e**). In these 48 hpi samples, bacteria were larger and displayed a relaxed genome, in contrast to the bacteria observed at 6 hpi that were smaller and had a more compact cytosol (**Fig. 1d**). These data are consistent with the *C. burnetii* inside these vacuoles transitioning from the SCV to LCV. Moreover, a large portion of intracellular bacteria observed at 48 hpi were undergoing cell division, indicating that they are metabolically active and in the LCV state.

Several bacteria in the LCV state were intimately associated with the CCV membrane at 48 hpi. The spacing between the bacterial outer membrane and vacuole membrane was ∼23.2 nm (**Extended Data Fig. 2c, d, f**). Closer examination of these contact sites revealed intermembrane tethers ∼18.2 nm long extending from the bacterial outer membrane to the vacuolar membrane (**Extended Data Fig. 2c, d, g)**. Numerous small vesicles were also visible inside the CCV at 48 hpi, and host microtubules were closely associated with the vacuolar membrane (**Fig. 1e, Extended Data Fig. 3**). Taken together, these results demonstrate that a combination of cryo-fluorescence microscopy, cryo-FIB milling, and cryo-ET imaging enables target-specific visualization of intracellular pathogens in their native host environment at unprecedented resolution.

### Distinct features of the LCV and SCV revealed by cryo-ET

To characterize the differences between *C. burnetii* LCVs and SCVs, bacterial cells were isolated from *C. burnetii*-infected host cells at 48 hpi and 168 hpi (**Fig. 2a-f**). Low magnification projection images were first acquired from these samples to estimate the composition of *C. burnetii* inside the CCV at these two time points (**Fig. 2a, d**). Bacteria were distributed into different categories based on size to determine the relative proportion of each variant in the population. The average LCV was ∼1,376 nm long and ∼507 nm wide, whereas the average SCV was ∼652 nm long and ∼275 nm wide (**Fig. 2h-i**). Therefore, the length and width of an LCV is reduced by roughly half upon transition from the LCV to SCV form. A population of transitional cells of intermediate size compared to the LCV and SCV forms was also observed. Samples from host cells infected for 48h contained 82% LCVs and 18% transitional cells. By contrast, samples isolated from cells infected for 168h contained 21.5% LCVs, 26.3% transitional cells, and 52.2% SCVs (**Fig. 2g**). This observation is consistent with the intercellular *C. burnetii* population shifting from LCVs to SCVs during this late stage of host cell infection.

**Figure 2.**
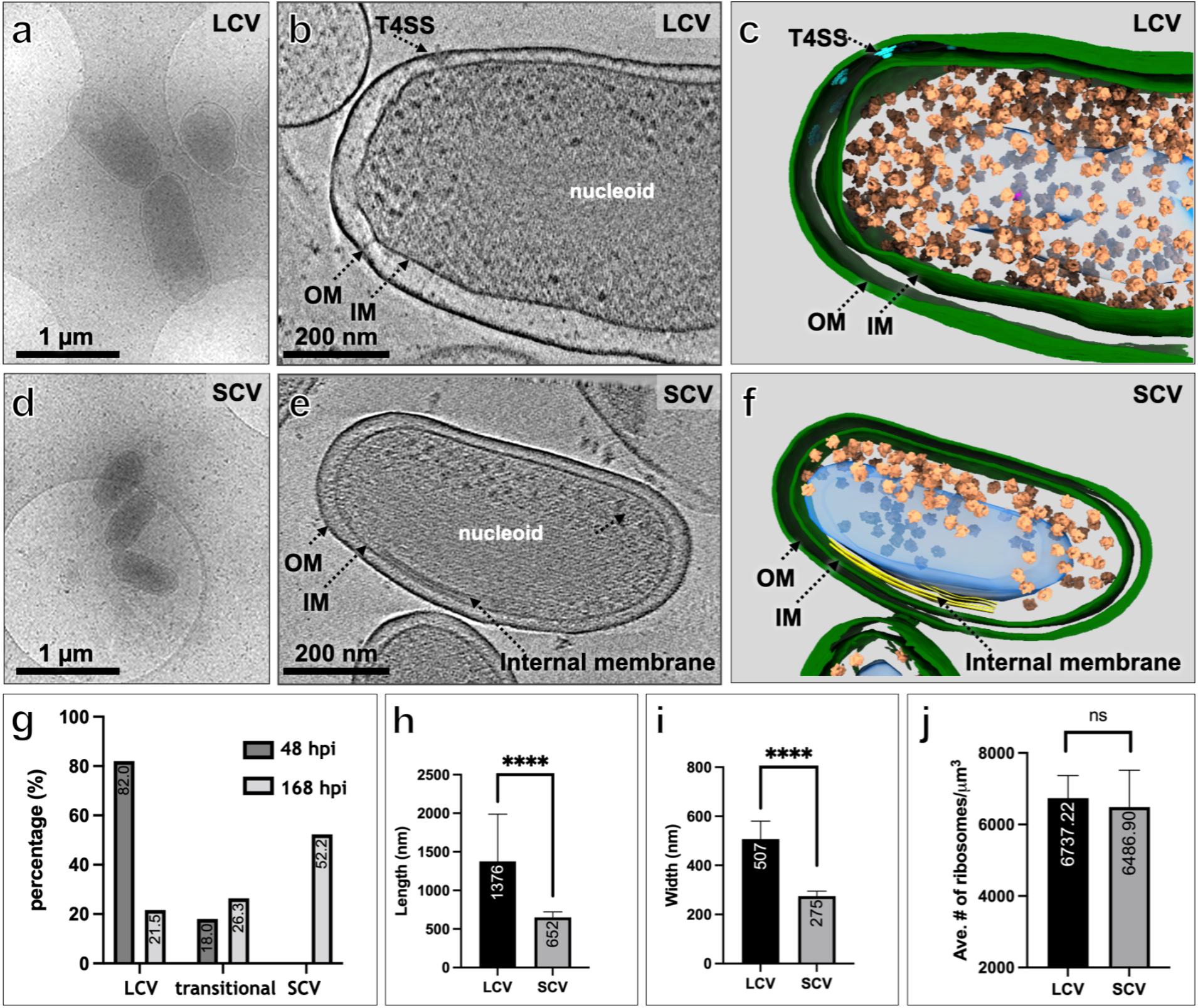
Visualization of *C. burnetii* undergoing significant morphological transformations. **(a)** A low-magnification cryo-EM image of large cell variants (LCVs), which were purified from 2-day-old CCVs. **(b, c)** A central tomographic section and a 3D view of a representative LCV show detailed structures of T4SS and ribosomes. **(d)** A low-magnification cryo-EM image of small cell variants (SCVs), which were purified from 7-day-old CCVs. **(e, f)** A central tomographic section and a 3D view of a representative SCV show detailed structures of internal membranes and ribosomes. (**g**) Quantifications of different developmental forms in 2-day-old (N=206) and 7-day-old (N=391) CCVs. **(h-i)** Comparisons of the length and width of the two variants, respectively. Ns: p-value > 0.05. ****: p-value < 0.001. (N=20). (**j**) Comparison of the number of ribosomes per cell volume in SCV (N=3) and LCV (N=2).

Cryo-ET imaging of the SCVs showed increased density in the periplasmic space compared to LCVs, likely the result of extensive cell wall remodeling leading to a thickening of the peptidoglycan layer in the SCVs^31^ (**Fig. 2b, e**). These images also showed that the LCV contains a more diffuse and loosely organized nucleoid compared to the densely packed nucleoid of SCVs (**Fig. 2b, e**). The densely packed nucleoid in the SCV likely represents the heterochromatic state of genomic material not actively being transcribed^32^. Ribosome positions from tomograms negatively correlate with the position of the nucleoid, as previously demonstrated in *Escherichia coli*^33^ (**Fig. 2b-c, 2e-f**). At various levels of densities, ribosomes were spread throughout the cytoplasm of the LCV, including the central regions (**Fig. 2b, c**), whereas ribosomes were exclusively found in one corner of the cytoplasm and strictly excluded from the nucleoid in the SCV, likely due to densely packed genomic material (**Fig. 2e, f**). Data from these tomograms indicate that the average LCV has 6,737 ribosomes/μm^3^, and the average SCV has 6,486 ribosomes/μm^3^, not a statistically significant difference (**Fig. 2j**). Thus, *C. burnetii* maintains a similar number of ribosomes per cell volume in both the LCV and SCV.

### Inner membrane preservation during *C. burnetii* developmental transitions

Intriguingly, imaging of SCVs revealed a stack of tightly packaged membranes next to the cytoplasmic face of the inner membrane (**Fig. 2e, f**). The presence of internal membranes with an unusual structurein *C. burnetii* was reported over four decades ago using TEM images of chemically fixed specimens^28^. To investigate the biogenesis of the membrane folds identified in the SCV, we closely examined inner membrane regions in tomograms of purified *C. burnetii* at different developmental stages. These images showed a gradual progression of folding of the inner membrane inward toward the cytosol as *C. burnetii* transitioned from an LCV to SCV. LCV images displayed a small stretch of inner membrane invagination (**Fig. 3a-c**). Transitional cells showed a longer stretch of inner membrane invagination and assembly of these membranes into folded structures (**Fig. 3d-f**). Transition to the SCV resulted in organization of these membranes into defined structures consisting of 2-6 tightly stacked membranes **(Fig. 3g-i**) with a distance between each layer of ∼7 nm (**Fig. 3j**). These membrane stacks were consistently located next to the condensed bacterial genome, and membrane tracing revealed that the stacks are contiguous with the inner membrane (**Fig. 3c, f, i**). These results suggest that packaging of the inner membrane takes place during contraction of the cell as it transitions from an LCV to a SCV. FIB-milling analysis of *C. burnetii* inside a CCV in infected host cells also revealed folding of the membrane (**Extended Data Fig. 4a-d**). At 48 hpi, similar sites of membrane invagination were visible in LCVs (**Extended Data Fig. 4c-d)**. Thus, the membrane invagination phenotype observed in the LCV imaged from isolated bacteria was also displayed in LCVs contained within the host cell vacuole. Intriguingly, at 6 hpi, a large round bubble could be observed directly beneath the inner membrane of *C. burnetii* presumably undergoing the SCV to LCV transition (**Extended Data Fig. 4a-b)**. Overall, these data suggest that the inner membrane stack can unfold to expand into LCV, presumably allowing rapid transitions between the two developmental forms (**Fig. 3k**).

**Figure 3.**
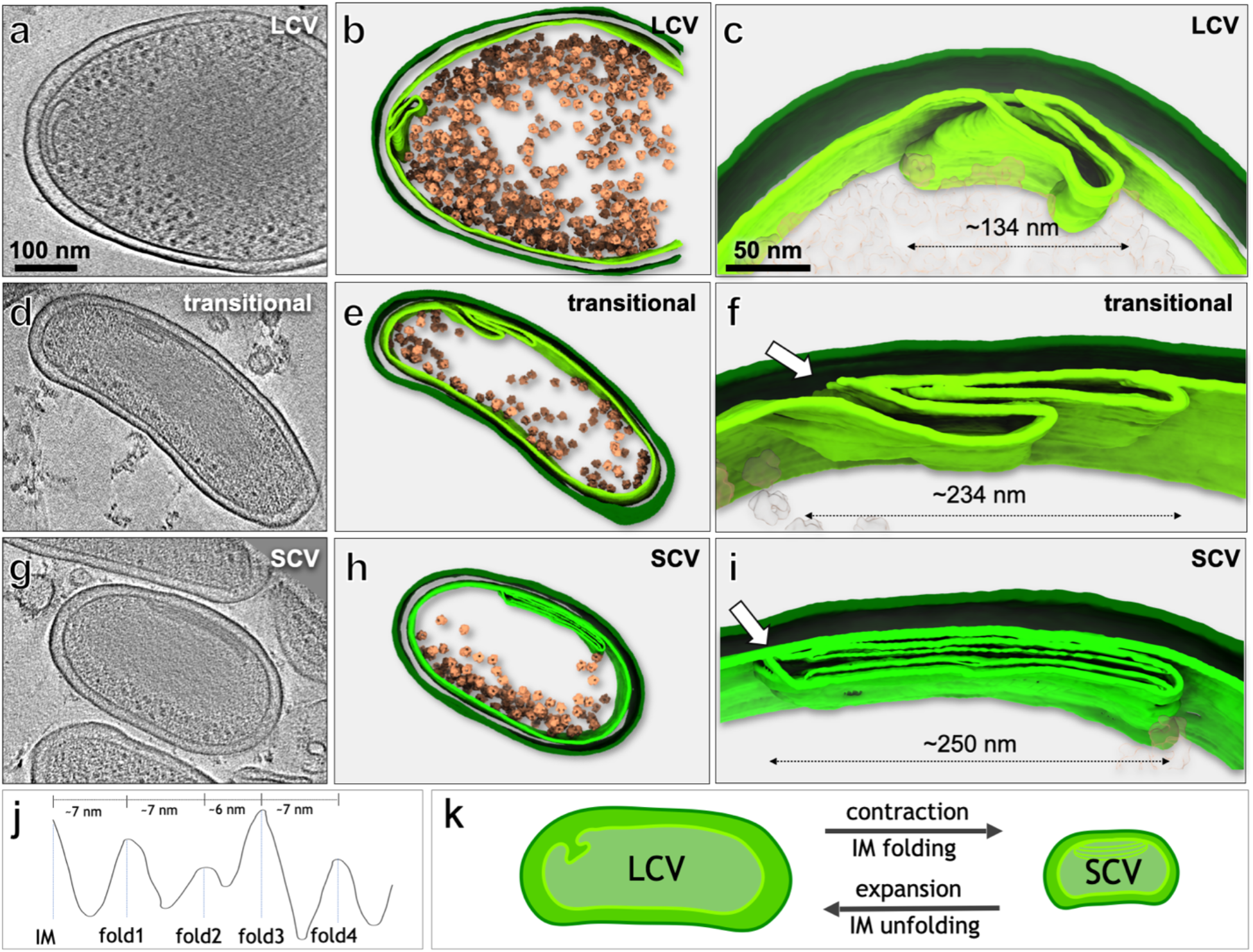
The inner membrane may be involved in the rapid contraction and growth of *C. burnetii*. **(a, b)** A central section and a 3D view of an LCV tomogram show additional membrane stacks in the cytoplasm. **(c)** A zoom-in view of the membrane stacks present in LCVs. **(d, e)** A central section and a 3D view of a transitional variant tomogram show extended membrane stacks in the cytoplasm. **(f)** A zoom-in view of the membrane stacks with a length of ∼234 nm. The junctions between the inner membrane and the cytoplasmic membrane stacks are indicated with white arrows. **(g, h)** A central section and a 3D view of an SCV tomogram show extended membrane stacks in the cytoplasm. **(i)** A zoom-in view of the membrane stacks with a length of ∼250 nm. **(j)** Graph measuring the membrane spacing within the membrane stacks shown in panel i. **(k)** Schematic model of membrane remodeling during *C. burnetii* developmental transitions.

### Assembly of the Dot/Icm machine correlates with *C. burnetii* developmental transition

Dot/Icm machines are required for the establishment of the CCV. Although genes encoding components of the machine are related to those in *L. pneumophila, C. burnetii* Dot/Icm machines have not been characterized in detail. Leveraging our cryo-ET reconstructions at different stages of *C. burnetii* developmental transitions, we had a unique opportunity to thoroughly analyze the Dot/Icm machines using approaches similar to those previously described in several studies of *L. pneumophila*^23-26^ (**Fig. 4a**). Intriguingly, Dot/Icm machines are present in LCVs replicating in host vacuoles (**Extended Data Fig. 5a-d**), while they are not visible in SCVs (**Fig. 4b**) or a *dotC* transposon mutant (**Fig. 4c**). Given that *dotC* encodes a core outer membrane component essential for the initial stage of Dot/Icm secretion machine assembly^25,34^, our data suggest that the assembly of Dot/Icm machines correlates with the developmental transition from SCV to LCV.

**Figure 4.**
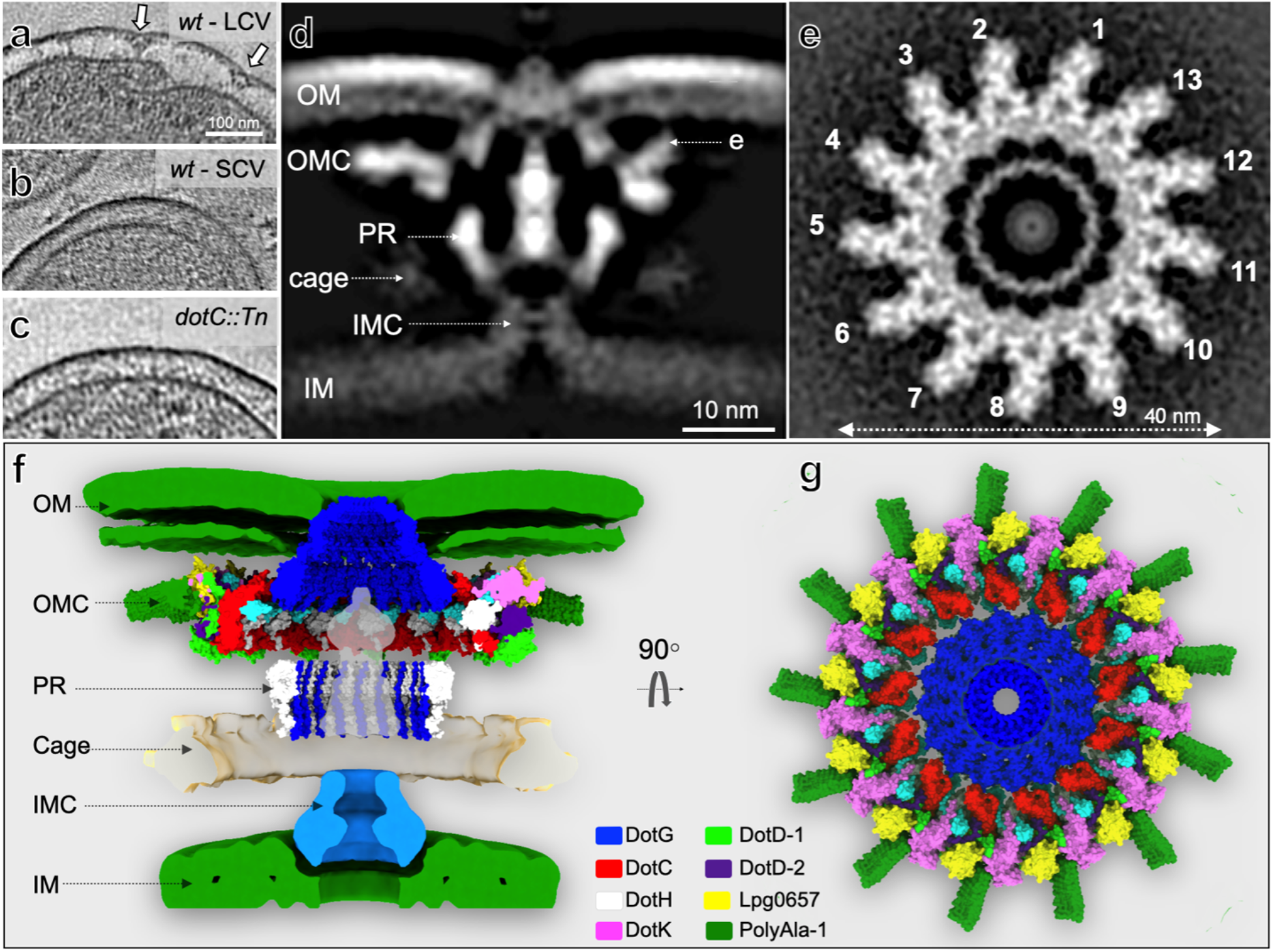
The molecular architecture of the *C. burnetii* T4SS. (**a-c**) Tomographic slices of representative wild-type (*wt*) LCV, SCV, and *dotC* transposon mutant (*dotC*::Tn) LCV. T4SSs are only visible in LCVs. **(d)** A central slice of the *in-situ* structure of the *C. burnetii* T4SS. IM: inner membrane, OM: outer membrane, PG: peptidoglycan, OMC: outer membrane complex, PR: periplasmic ring, IMC: inner membrane complex. **(e)** A cross section of the OMC at the position indicated in panel **c**. (**f, g**) A side view and a top view of the model of the *C. burnetii* T4SS.

To determine an *in-situ* structure of the *C. burnetii* Dot/Icm machine, we used subtomogram averaging to analyze 7,905 particles from transitional cells and LCVs. This resulted in a*C. burnetii* Dot/Icm machine map that revealed a structure similar to that of *L. pneumophila* (**Fig. 4d-e**). Both machines have highly conserved components: outer membrane complex, periplasmic complex, inner membrane complex, and cage (**Fig. 4d-e, Extended Data Fig. 6a-b**). Specifically, the outer membrane complex is composed of 13 spokes with an overall width of ∼40 nm (**Fig. 4e, Extended Data Fig. 6c-d**). Each spoke was further refined and compared with the near-atomic structure of the outer membrane complex derived from the *L. pneumophila* Dot/Icm machine^21,22^ (**Extended Data Fig. 7a-e**).

The *C. burnetii* Dot/Icm system displays distinct features compared to *L. pneumophila*^24,26^. The outer membrane complex of *C. burnetii* was relatively flat except for a pinching inward around the outer membrane penetrating the DotG complex (**Fig. 4 f-g, Extended Data Fig. 6a, b**). The Dot/Icm outer membrane complex assembles around the flat membrane with a ∼5 nm space between the two. The rotation and flattening of the outer membrane complex may result from assembly along the flat outer membrane. Moreover, the *C. burnetii* Dot/Icm system displays an A-shaped inner membrane complex, unlike the V-shaped inner membrane complex observed in *L. pneumophila*^23-26^ (**Extended Data Fig. 6g-h**). The inner membrane complex is predicted to translocate effector molecules from the bacterial cytosol across the inner membrane. Raw tomograms identified a subpopulation of Dot/Icm machines that lack the density corresponding to the inner membrane complex.

To examine the heterogeneity of inner membrane complex assembly, we performed classification analysis around this complex. The results indicate that 38.6% of the Dot/Icm machines identified from LCV images displayed an assembled inner membrane complex. By contrast, none of the bacteria in the transitional phase displayed a Dot/Icm-associated inner membrane complex, indicating that the complete core complex assembles upon transition to the LCV form (**Fig. 5**). Overall, this cryo-ET study predicts that *C. burnetii* achieves rapid developmental transitions through a process involving preservation of the inner membrane and that this transition corresponds with the assembly of a functional Dot/Icm machine critical for intracellular survival (**Fig. 5, Supplementary Video 2**).

**Figure 5.**
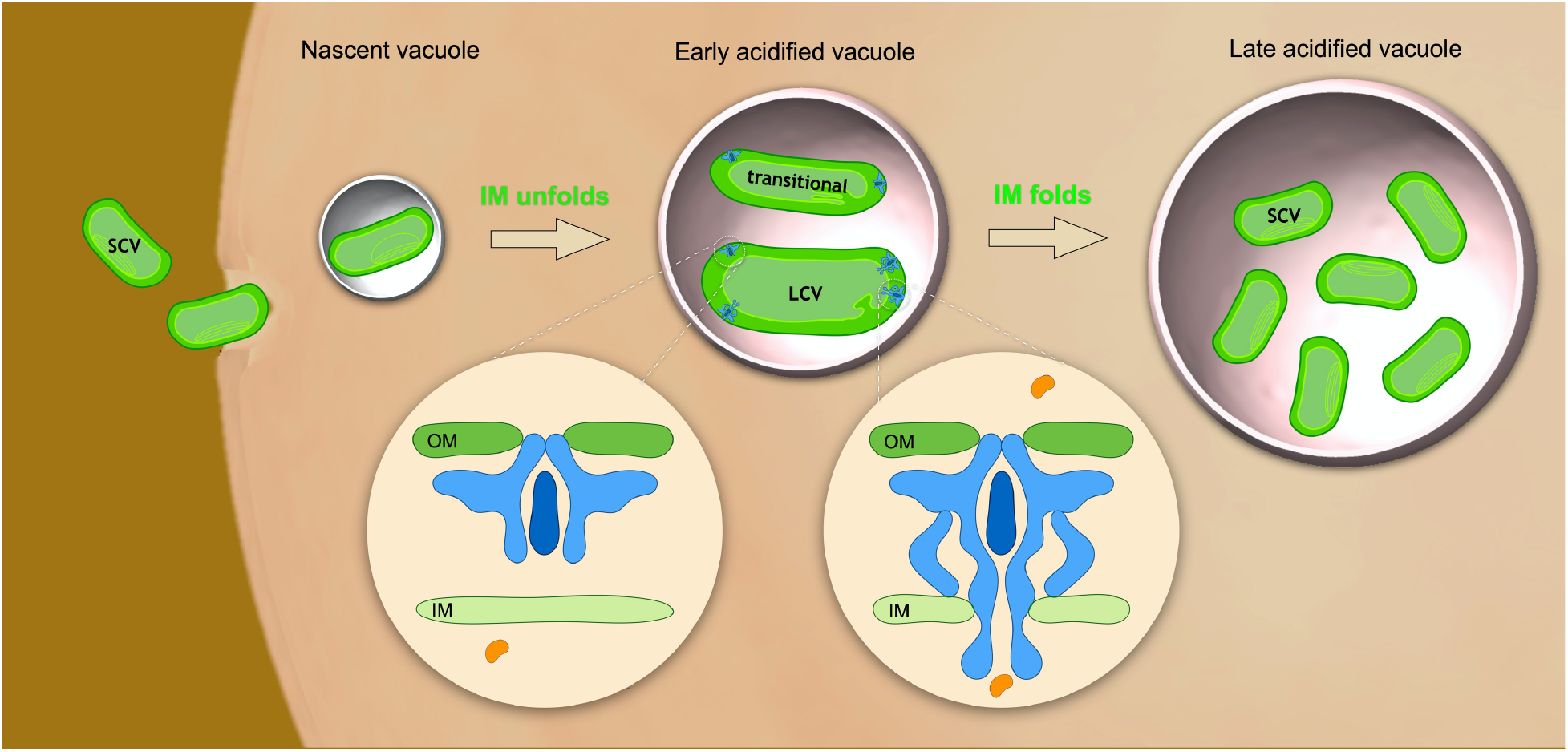
Model of how developmental transitions enable T4SS assembly in *C. burnetii*.

Another striking feature of *C. burnetii* is the outer membrane, which has an outer leaflet significantly thicker (∼3.5 nm) than the inner leaflet (∼1.2 nm) (**Extended Data Fig. 6e**). We previously demonstrated that the outer membrane of *L. pneumophila* is composed of an extra layer of electron density on top of the outer leaflet that possibly consists of LPS or surface proteins^24^. The outer membrane of *C. burnetii* does not show this extra layer (**Extended Data Fig. 6e, f)**.

## DISCUSSION

*C. burnetii* cells at different developmental stages have been visualized *in situ* using an advanced cryo-ET pipeline that enables high-resolution views of cells and their highly specialized Dot/Icm machine. Subtomogram averaging and classification analysis of the Dot/Icm machines led to the conclusion that a complete core complex assembles as bacteria transition from the infectious SCV to the replicative LCV form. Multiple studies in *L. pneumophila* have shown that the Dot/Icm system assembles at cell division sites during bacterial replication and remains localized primarily at the poles of the bacilli^23-26,35,36^. Most of the “WiFi-like” structures representing an assembled Dot/Icm machine in *C. burnetii* were identified at the poles of LCV bacilli, suggesting a similar Dot/Icm machine assembly process in both *C. burnetii* and *L. pneumophila*.

The absence of assembled Dot/Icm machines in SCV bacteria raises the important question of when a functional Dot/Icm machine is needed for host infection by *C. burnetii*. These data reinforce studies showing that effector proteins are not translocated by *C. burnetii* during early stages of host cell infection and that effector translocation requires transport of *C. burnetii* to an acidified lysosome-derived organelle^37^. This niche-specific translocation ability suggests that endocytic transport of *C. burnetii* to a lysosome-derived organelle is sufficient to initiate the transition from SCV to LCV and to promote bacterial cell division. Subsequent expression and production of Dot and Icm proteins would then result in the assembly of a Dot/Icm machine that can deliver effector proteins into the host cell. Consistent with this hypothesis, peptides generated in the lumen of the hydrolytic lysosome-derived vacuole in which *C. burnetii* resides are sensed by the bacterial PmrAB two-component regulatory system^38^, which regulates many of the genes encoding Dot and Icm proteins^39,40^. Thus, the Dot/Icm system plays an important role in modulation of host cell functions once *C. burnetii* replication has been initiated, and this role likely involves suppressing host defenses that limit bacterial replication and modifying the vacuole to ensure sufficient nutrient availability for bacterial replication.

Intermembrane tethers were observed between the vacuolar membrane and the outer membrane of *C. burnetii* LCVs when *C. burnetii*-infected host cells were examined at 48 hpi (**Extended Data Fig. 2c, d, g**). Similar tethers were reported in a TEM image from *C. burnetii*-infected cells^41^ in a study that had speculated that these tethers represent regions where the Dot/Icm machine has engaged the membrane of the CCV. Our data do not support this hypothesis as the location of the tethers does not correlate with the location of Dot/Icm machines. Nonetheless, our data do not rule out the possibility that the formation of these tethers is stimulated by the activity of the Dot/Icm machine. It is also possible that the intermembrane tethers represent interaction sites between bacterial adherence factors and the vacuolar membrane that facilitate translocation of Dot/Icm effectors into the host cell by stabilizing intimate contacts. Recently, similar cryo-FIB milling imaging of *L. pneumophila* inside amoeba hosts showed that the Dot/Icm machine mediates interaction with the vacuolar membrane to form a membrane indentation that facilitates effector secretion^36^. However, we did not observe such a phenomenon between the CCV membrane and *C. burnetii* secretion machines. It is unlikely that these observations indicate fundamental differences between Dot/Icm machines in *L. pneumophila* and *C. burnetii*. The differences observed may reflect the type of host (i.e. amoeba vs. mammalian cells) and the mechanism each bacterium uses to create replication vacuoles for survival in specific niches.

Our extensive analysis of the morphology of *C. burnetii* revealed a population of intermediate-sized cells, called transitional variants, that seem to be an intermediate form between LCVs and SCVs. Transitional variants were as narrow in cell width as SCVs but had a cell length similar to that of LCVs. A bacterial cell elongation model, whereby bacilli maintain a constant width while increasing in length, has been described in detail for many Gram-negative bacteria, including *Escherichia coli* and *Caulobacter crescentus*^*42,43*^. The transition from SCV to LCV involves a change in both cell width and length. It is not well understood how this dramatic change in cell morphology occurs in *C. burnetii*. The observation that transitional variants display the width of SCVs and length of LCVs suggests that, upon entry into host cells, the infectious SCV cells first expand in length before becoming wider. A study that monitored changes in gene expression levels of *C. burnetii* at different time points after infection showed that the expression of genes encoding rod shape-determining proteins such as RodA, MreB, and MreC was downregulated in SCVs^31^. These proteins regulate synthesis of the peptidoglycan cell wall, and thus their differential expression may coordinate dynamic transitions in cell shape. Additional studies are needed to determine how these proteins in *C. burnetii* contribute to cell shape and regulate the SCV-to-LCV-to-SCV cycle inside host cells.

Our population analysis of *C. burnetii* purified from 48 hpi and 168 hpi CCVs (**Fig. 2g**) indicates that transitional variants account for a small fraction of the morphological forms of *C. burnetii* at both time points. This result suggests that the transition between SCV and LCV occurs relatively quickly. We observed a developmentally regulated pathway that results in the assembly of inner membrane folds during the transition from LCV to SCV. We propose a model whereby preservation of the bacterial membrane as the *C. burnetii* cell contracts during the LCV-to-SCV transition facilitates rapid cell elongation after the dormant SCV form infects a new host cell and transitions back to LCV form (**Fig. 3k**). The high-resolution cryo-ET imaging presented here provides evidence that an unusual internal membrane structure first reported in *C. burnetii* decades ago^28^ is involved in developmental transitions. Future studies may provide the molecular mechanism underlying how the inner membrane stacks are generated and how they contribute to cell differentiation and this unorthodox mechanism of cell growth.

## METHODS

### Bacterial growth conditions

*C. burnetii* Nine Mile phase II (NMII), strain RSA439 (clone 4)^44^, and mCherry-expressing transposon mutants^45^ were cultured axenically in liquid acidified citrate cysteine medium 2 (ACCM-2) for 2-7 days at 37°C, 5% CO_2_ and 2.5% O_2_, as previously described^29,46^. *C. burnetii* strains used in this study are listed in **Table 1**. A strain having a transposon insertion in a neutral intergenic region of the chromosome (ig::Tn) was previously shown to display intracellular replication comparable to the parental *C. burnetii* NMII strain^47^. When appropriate, chloramphenicol (3 µg/ml) was added to ACCM-2. To calculate MOIs, *C. burnetii* genome equivalents (GE) were determined by quantitative PCR (qPCR) using *dotA*-specific primers as previously described^45^.

**Table 1.**
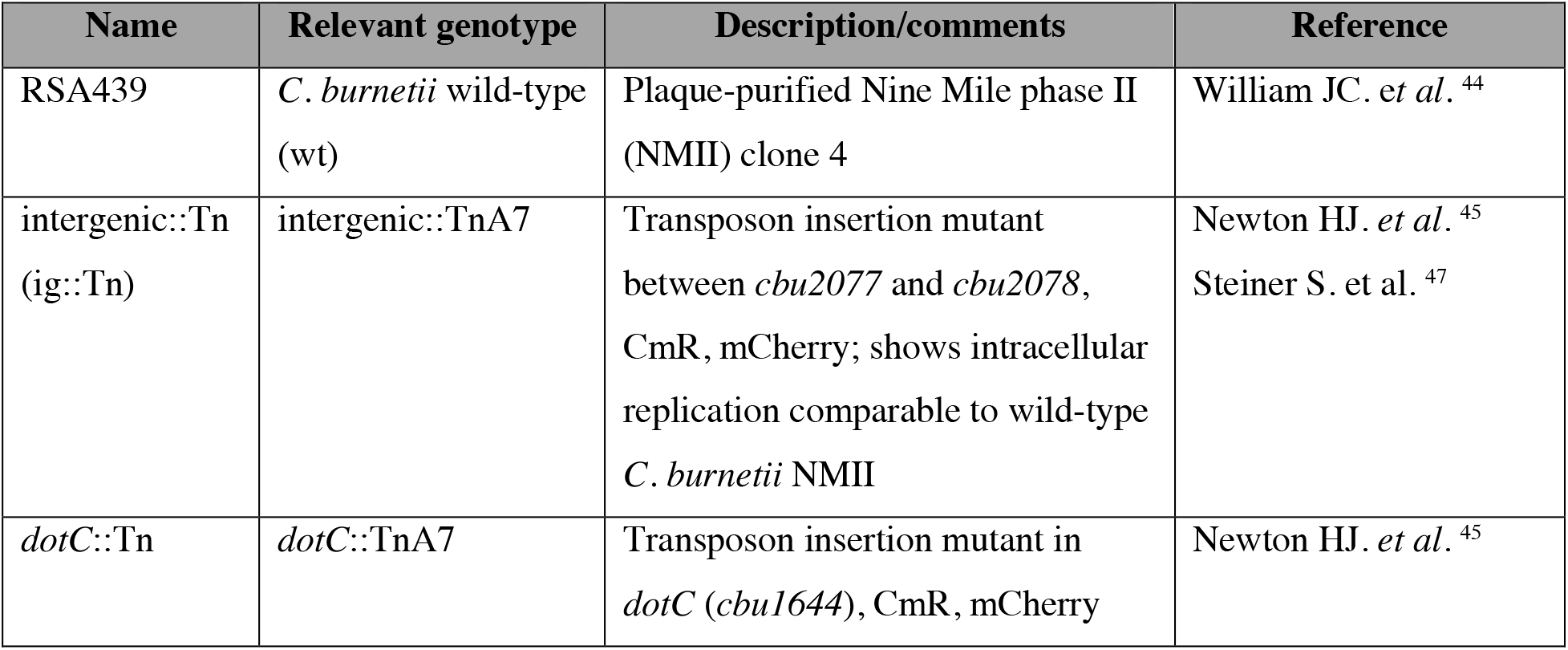
Bacterial strains used in this study.

Extended Data
| Name | Relevant genotype | Description/comments | Reference |
| --- | --- | --- | --- |
| RSA439 | *C. burnetii* wild-type (wt) | Plaque-purified Nine Mile phase II (NMII) clone 4 | William JC. *et al.* <sup>44</sup> |
| intergenic::Tn (ig::Tn) | intergenic::TnA7 | Transposon insertion mutant between *cbu2077* and *cbu2078*, CmR, mCherry; shows intracellular replication comparable to wild-type *C. burnetii* NMII | Newton HJ. *et al.* <sup>45</sup><br>Steiner S. *et al.* <sup>47</sup> |
| *dotC*::Tn | *dotC*::TnA7 | Transposon insertion mutant in *dotC* (*cbu1644*), CmR, mCherry | Newton HJ. *et al.* <sup>45</sup> |

**Table 2.**
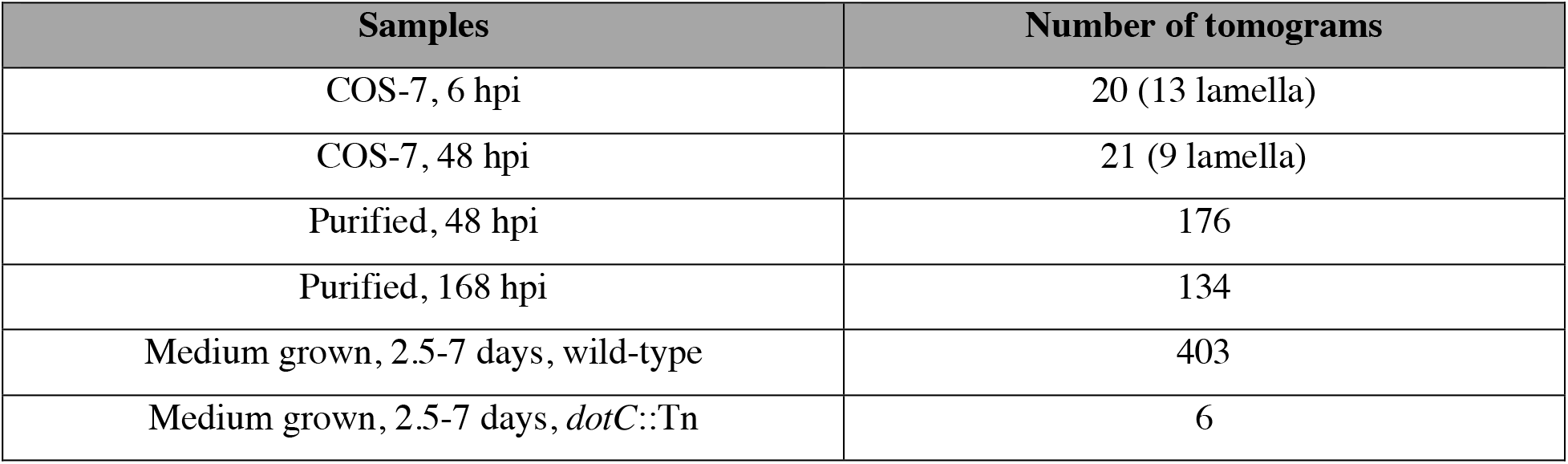
Summary of cryo-ET data used in this study.

### COS-7 cell culture on cryo-EM grids and *C. burnetii* infection for FIB-milling experiments

COS-7 cells were cultured in DMEM supplemented with 10% fetal bovine serum (FBS) at 37°C/5% CO_2_. Prior to seeding COS-7 cells on the cryo-EM grids, gold cryo-EM grids with R1/4 or R2/4 Quantifoil were coated with 0.1 mg/ml poly-D-lysine for 1 hour, rinsed 4 times with sterile water, and incubated with the fully supplemented medium at 37°C/5% CO_2_ for >8 hours. Freshly trypsinized COS-7 cells were seeded on the pre-treated EM grids and allowed to grow overnight at 37°C/5% CO_2_.

To observe CCVs at different time points, a *C. burnetii* strain producing the mCherry protein was used. To analyze CCVs at 6 hpi, adherent COS-7 cells on EM grids were infected at an MOI of 300 in DMEM with 10% FBS. Cells were washed with PBS at 4 hpi and fresh cell culture medium was added. Infection was carried out for 6 additional hours. To analyze CCVs at 48 hpi, experimental procedures were as above, but infection was performed at an MOI of 100 and was carried out for a total of 48 hours after washing cells with PBS.

### Isolation of intracellular *C. burnetii*

To purify intracellular *C. burnetii*, COS-7 cells were cultured in a T-75 flask at 50% confluency and infected at an MOI of 100. COS-7 cells were washed with PBS at 4 hpi, fresh medium was added to cells, and infection was carried out for 48 or 168 hours. The medium was removed, and 20 ml of PBS was added to the flask. Infected cells were harvested using a cell scraper and the cell suspension was pipetted several times and vortexed to break CCVs and release *C. burnetii* into the supernatant. The cell suspension was spun at 1,000 rpm/4°C for 10 minutes to pellet large cellular debris from COS-7 cells. The supernatant was filtered through 1 layer of Kimwipe to further remove cellular debris.

### Cryo-ET sample preparation

Bacterial cultures were mixed with 10-nm colloidal gold particles at 4:1 ratio for the marker-dependent tilt series alignment. The mixture was deposited onto freshly glow-discharged copper grids with R2/1 Quantifoil for 1min, blotted with filter paper, and frozen in liquid ethane using a gravity-driven homemade plunger apparatus.

For FIB-milling samples, DMEM supplemented with 10% FBS and 10% glycerol was added to the grids containing infected COS-7 cells for ∼1 minute to prevent the formation of crystalline ice inside COS-7 cells prior to plunge freezing in liquid ethane as described above.

### Cryo-fluorescence microscopy

Vitrified specimens were first clipped with autogrids designed for FIB-milling. Samples were loaded to the cryo-CLEM (Leica) microscope to image fluorescent *C. burnetii*. 6 × 6 Z-stacks, which cover ∼2/3 of the grid surface, were collected with bright field and mCherry channels. Projection images compatible with MAPS software were generated to correlate the fluorescence signals during the FIB-milling.

### FIB-milling of infected mammalian cells

FIB-milling was performed using an Aquilos FIB/SEM instrument. In order to correlate the cryo-fluorescence microscopy image with SEM image, a low-magnification montage of the entire grid was generated, and identifiable features were used to overlay the cryo-fluorescence image using the MAPS software. Samples were sputtered with a metallic platinum for 15 seconds followed by a coating with a layer of organometallic platinum (Pt) for 8 seconds to protect the sample. Additionally, 5 seconds of sputtering was done to prevent drifting during the milling. Samples were milled with a gallium ion beam to generate lamellae with <∼150nm thickness. Lastly, the lamellae were coated with 3 seconds of sputtering to reduce the charging effect when imaged with Volta phase plate (VPP).

### Cryo-ET data collection and reconstruction

Bacterial samples were imaged with a Titan Krios microscope (Thermo Fisher Scientific) equipped with a field emission gun, energy filter, VPP, and direct detection device (Gatan K2 Summit). VPP was used for the data acquisition at ∼1 µm defocus to increase the image contrast. The tomographic package SerialEM was used to collect 35 image stacks at a range of tilt angles between +51° to -51° (3° step size) using the bi-directional scheme with a cumulative dose of ∼60 e^-^/Å^2^. The magnification resulting in a pixel size of 2.75 Å at the specimen level was used. In addition, some bacterial samples were imaged with a Glacios microscope (Thermo Fisher Scientific) equipped with a direct detection device (Gatan K2 Summit) at 5 µm defocus using the same data acquisition strategy as described earlier. Each image stack containing 10-15 images were aligned using Motioncor2^48^ and then were assembled into the drift-corrected stacks by TOMOAUTO^49^. The drift-corrected stacks were aligned and reconstructed by IMOD marker-dependent alignment^50^.

Cryo-lamellae were imaged using the same Titan Krios microscope that had been upgraded with Gatan K3 camera. Similar data collection strategy was used as described in the previous paragraph. Tilt angle range was adjusted to account for the milling angle. The magnification resulting in a pixel size of 4.5 Å at the specimen level was used. Each image stack containing 10-15 images were aligned using Motioncor2^48^ and then assembled into the drift-corrected stacks by TOMOAUTO^49^. The drift-corrected stacks were aligned and reconstructed by IMOD^50^ marker-dependent alignment using the Pt residues as fiducials.

### Subtomogram averaging

I3 (0.9.9.3) was used for subtomogram analysis^51^. 7,905 T4SS machines from 713 tomograms were manually identified and extracted, as described previously^26^. 4×4×4 binned subtomograms were used for classification and junk-particle deletion. After a few rounds of alignment and classification, the OMC showed 13-fold symmetry; therefore, a 13-fold symmetry was imposed to assist the alignment process. Original subtomograms were used for the final structures.

Ribosomes shown in the segmentations were identified by the reference-based picking tool, averaged to ∼40 Å resolution, and mapped back into the original tomograms by EMAN 2.9^52^. Number of identified ribosomes inside SCVs and LCVs was used to calculate ribosome densities shown in **Fig. 2j**.

### Resolution estimation

The resolution estimation package ResMap^53^ was used to calculate local resolution of the focused refined outer membrane complex electron density map. The single-volume input method was used.

### 3D visualization

Many cellular features were segmented using the EMAN 2.9 segmentation tool^52^. Gaps in the membranes and features that were not successfully segmented by EMAN 2.9 were manually segmented using the Amira software. UCSF ChimeraX was used to visualize the segmentations and subtomogram average structures ^54^. PDB 6×62 was mapped into the focused refined map (**Extended Data Fig. 7c)** using the *Fitmap* command in UCSF ChimeraX. For the architectural model building, we applied C13 symmetry to PDB 6×62. PDB 6×64 was mapped into the corresponding periplasmic ring density using UCSF ChimeraX. Atomic model of DotG homolog was not available. We used *C. burnetii* DotG sequence (FASTA Q83B86-1) to predict the structure using AlphaFold2^55^. Atomic model predicted by the AlphaFold2 was mapped using a C16 symmetry as demonstrated by *L. pneumophila* DotG structure^22^.

## ACKNOWLEDGMENTS

We thank Jennifer Aronson for critical reading of the manuscript. We thank Shenping Wu for assisting with cryo-ET data collection. This work was funded by the National Institutes of Health (NIH) grants R01AI152421 to Jun Liu and Craig Roy and R01AI114760 to Craig Roy. Samuel Steiner was supported by the Early and Advanced Postdoc Mobility fellowships (P2BSP3_155237 and P300PA_164710) from the Swiss National Science Foundation (SNSF).

## Extended Data

**Extended Data Fig. 1.**
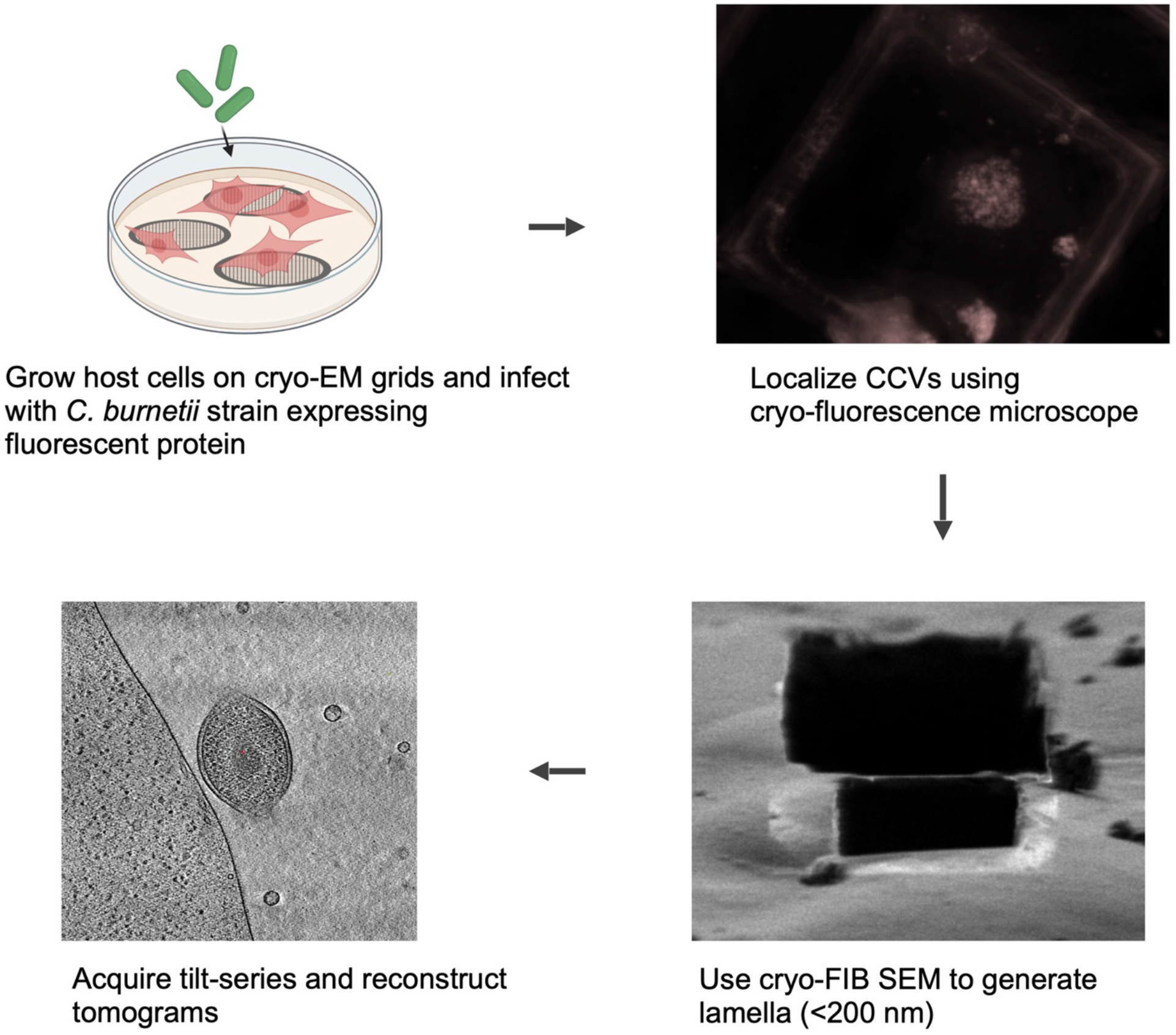
Summary of the experimental procedure to visualize an intracellular bacterial pathogen using cryo-fluorescence microscopy and FIB-milling.

**Extended Data Fig. 2.**
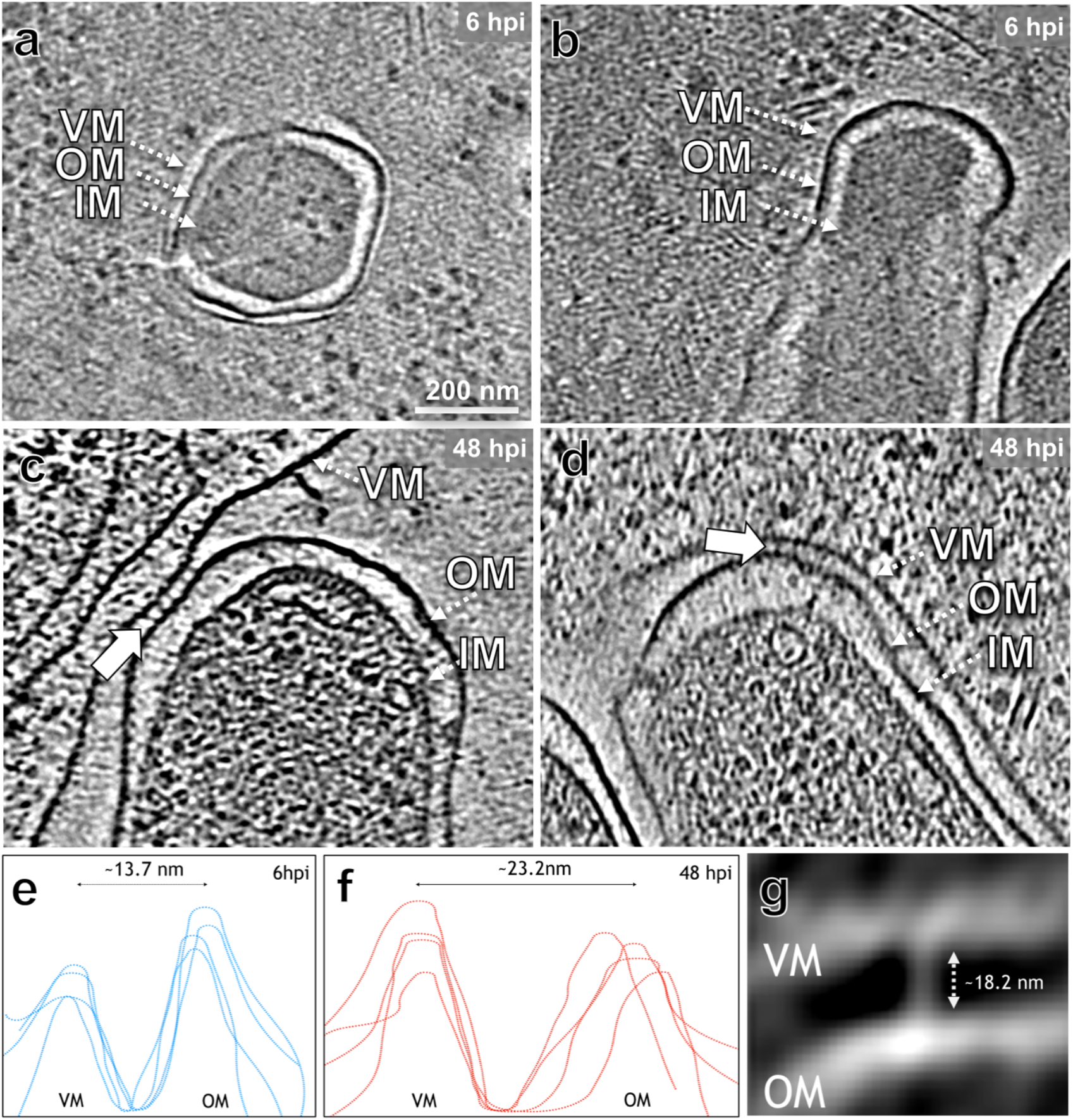
Membrane spacing between *C. burnetii* and the vacuolar membrane (VM) at different time points. (**a-b**) Tomographic slices show *C. burnetii* tightly surrounded by the CCV membrane at 6 hpi. IM: inner membrane, OM: outer membrane. (**c-d**) Tomographic slices show *C. burnetii* making close contact with the CCV membrane at 48 hpi. Blue arrows highlight the intermembrane tethers. (**e-f**) Membrane spacings between VM and OM at 6 hpi and 48 hpi, respectively, measured by IMOD. Peaks represent electron densities of membranes. (**g**) Subtomogram average structure of the intermembrane tethers found in between the VM and OM at 48 hpi.

**Extended Data Fig. 3.**
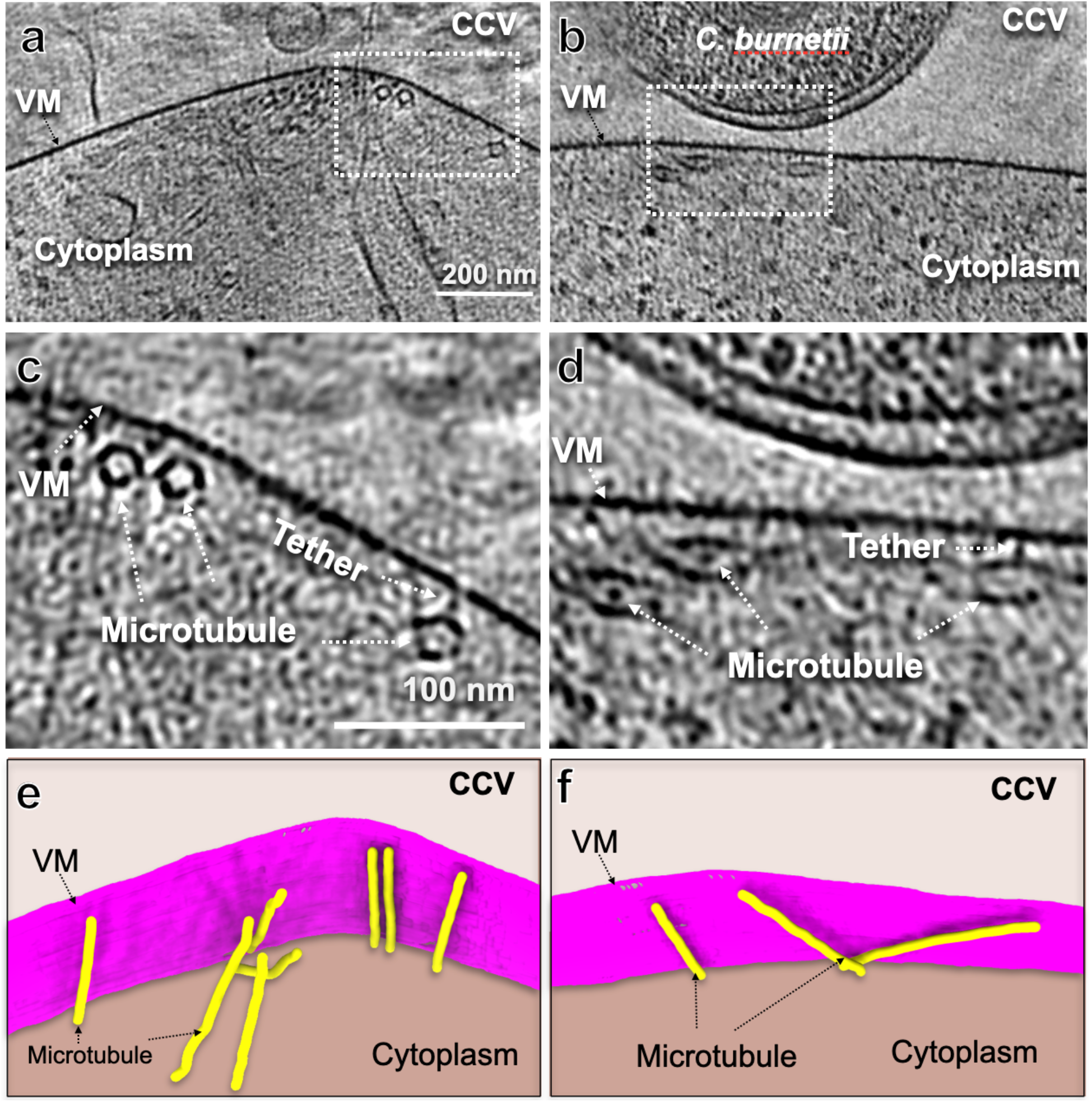
Microtubules make close contact with the CCV at 48 hpi. (**a, b**) Tomographic slices show the vacuolar membrane (VM) at 48 hpi. Many microtubules are closely associated with the VM. (**c, d**) Zoom-in views of highlighted boxes in panels a and b, respectively. The electron densities between microtubules and VM are highlighted with white arrows. (**e, f**) 3D renderings of tomograms in panels **a** and **b**, respectively.

**Extended Data Fig. 4.**
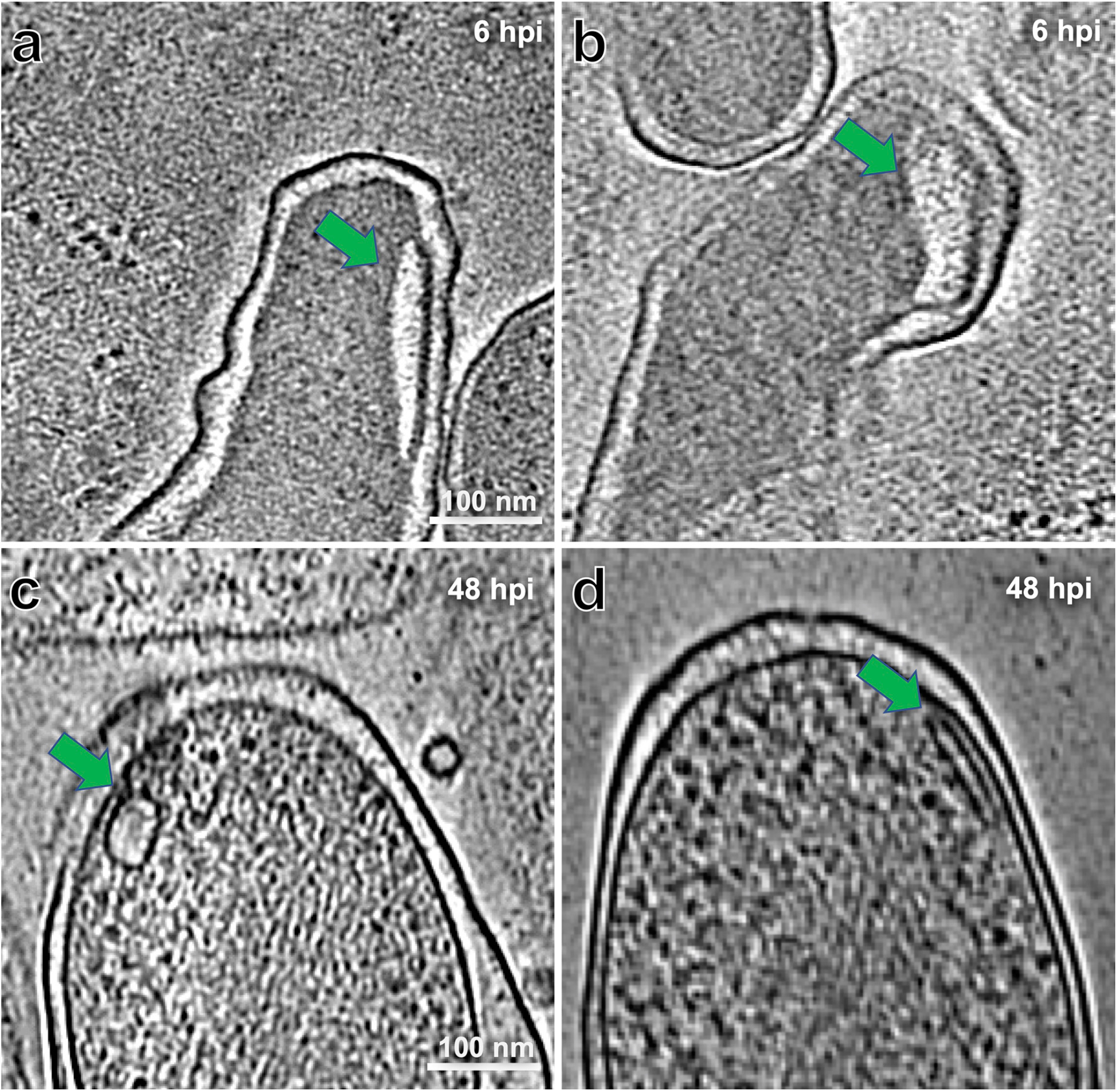
Membrane folding phenotypes found in intracellular *C. burnetii*. (**a, b**) Tomographic slices show *C. burnetii* at 6 hpi. Green arrows highlight the folded membrane. (**c, d**) Tomographic slices show *C. burnetii* at 48 hpi. Green arrows highlight the folded membrane, which is also found in purified *C. burnetii* at 48 hpi.

**Extended Data Fig. 5.**
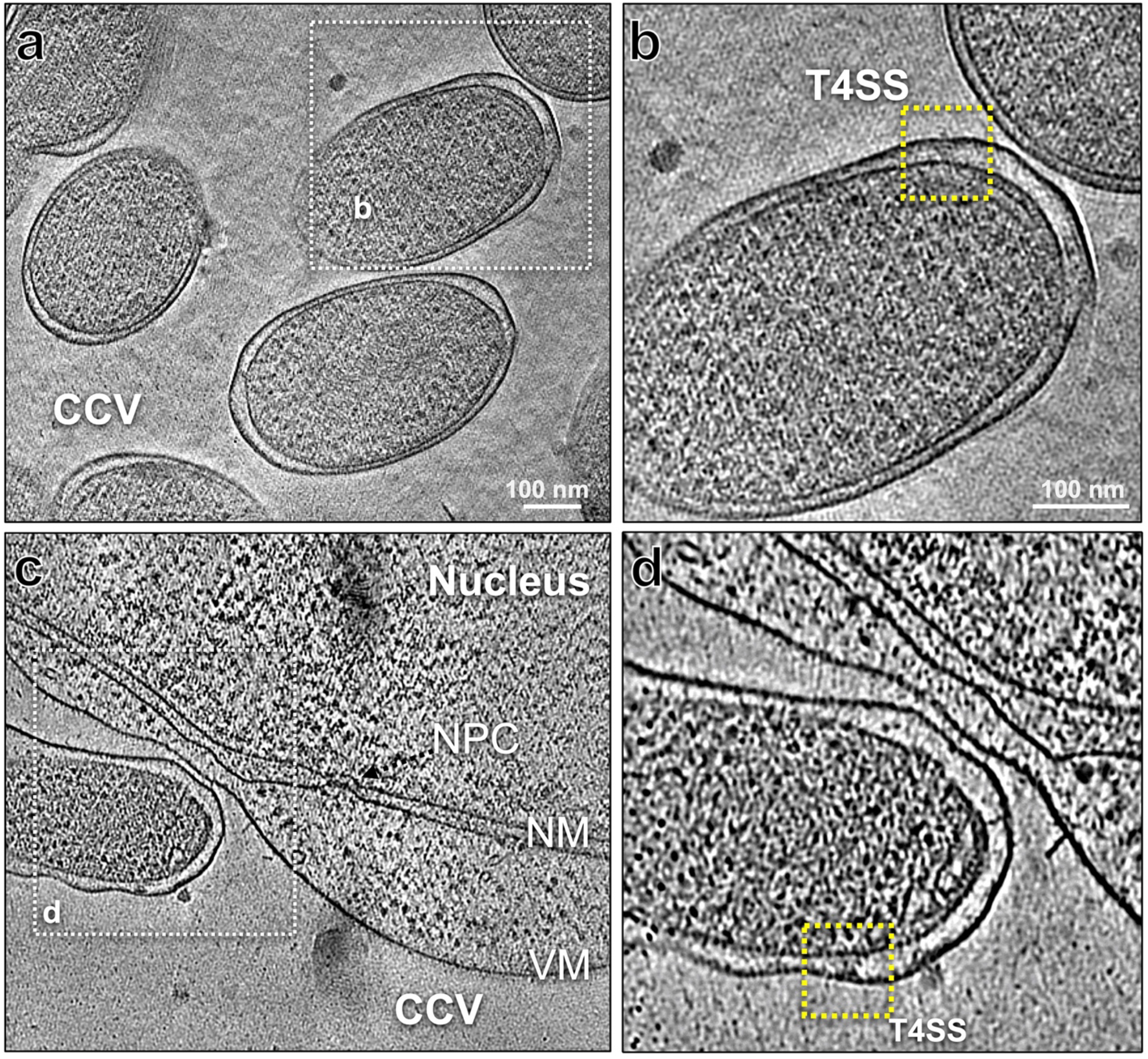
T4SS found in intracellular *C. burnetii* at 48 hpi. (**a, c**) Tomographic slices show *C. burnetii* inside a host cell. Panel **a** – in the middle of the CCV, panel **c** – near the vacuolar membrane (VM). (**b, d**) Zoom-in views of highlighted areas in panels a and c, respectively. T4SSs are boxed and labeled. NM: nuclear membrane, NPC: nuclear pore complex.

**Extended Data Fig. 6.**
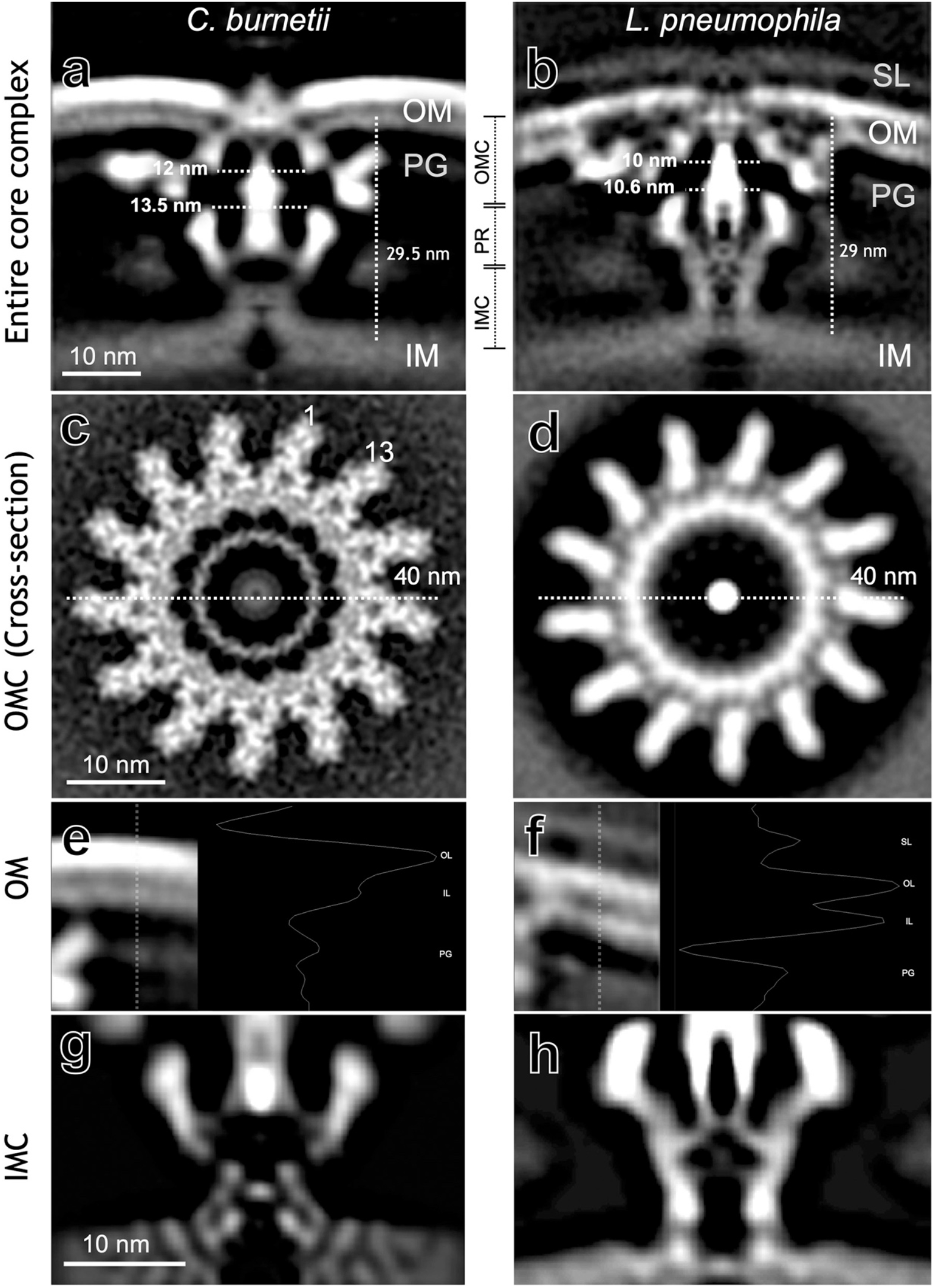
Comparison of *C. burnetii* and *L. pneumophila* T4SS. (**a-h**) Subtomogram average structures of *C. burnetii* and *L. pneumophila* T4SS core in side views (**a-b**), cross sections of the outer membrane complex (OMC) (**c, d**), side views of the outer membrane (OM) (**e-f**) and inner membrane complex (IMC) (**g, h**).

**Extended Data Fig. 7.**
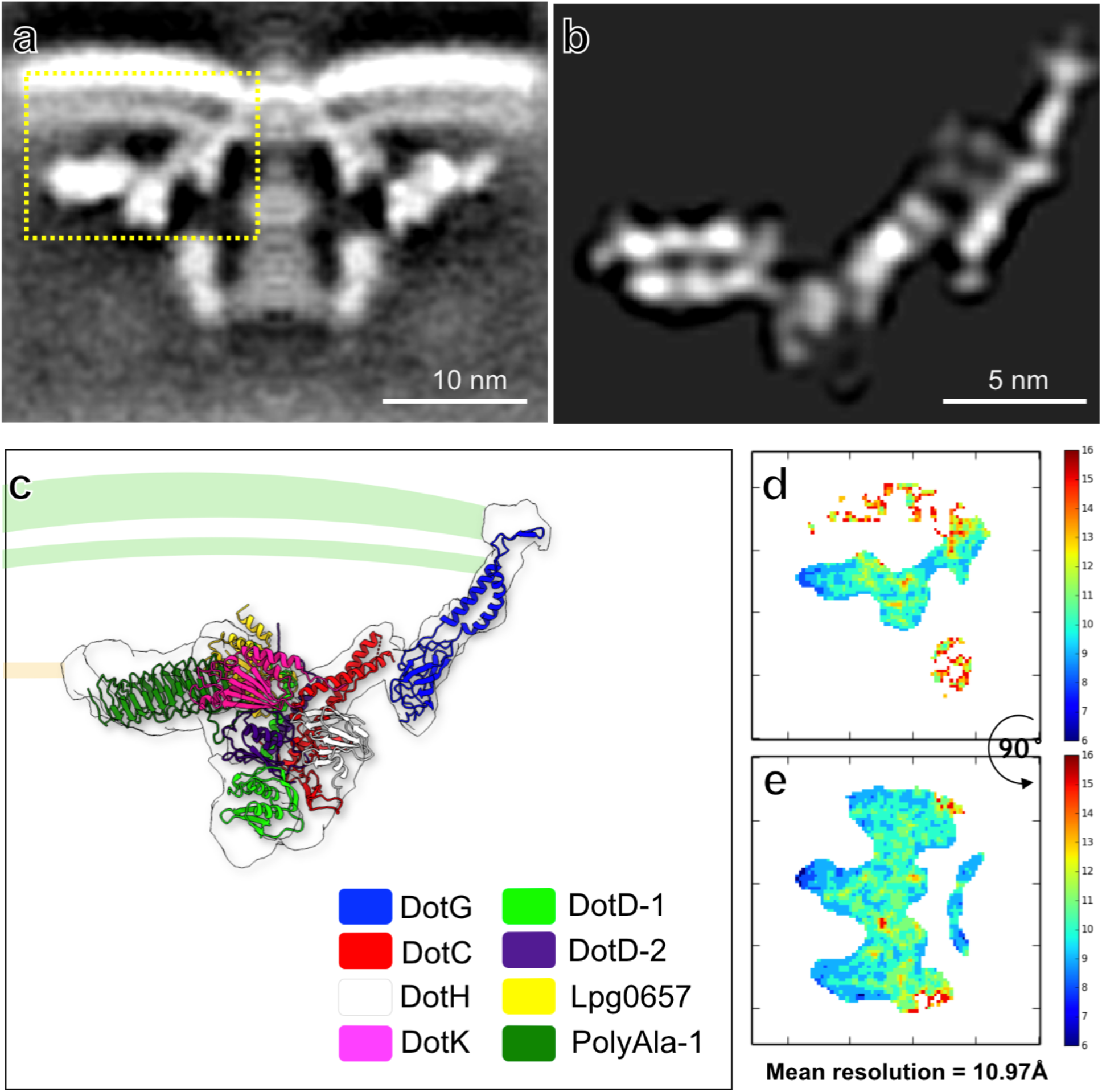
Focused refinement and model building of the *C. burnetii* T4SS. **(a)** 2D slice view of the subtomogram average structure of the *C. burnetii* T4SS. **(b)** Focused refined structure of the highlighted region in panel **a**. **(c)** Rigid fitting of PDB 6×62 into the density of the focused refined map. *C. burnetii* DotG was generated using AlphaFold2. (**d-e**) Local resolution estimated by ResMap software.

